# Corilagin controls post-parasiticide schistosome egg-induced liver fibrosis by inhibiting Stat6 signalling pathway

**DOI:** 10.1101/340299

**Authors:** Peng Du, Qian Ma, Jun Xiong, Yao Wang, Fan Yang, Feng Jin, Yun-Fei Chen, Zhen-Zhong Shang, Zhi-Lin Chen, Xuan Zhou, Hua-Rong Li, Lei Zhao

## Abstract

This study aims to explore the effect of Corilagin (Cor) on post-parasiticide schistosome egg-induced hepatic fibrosis through the Stat6 signalling pathway *in vitro* and *in vivo*. Cellular and animal models were established and treated by Corilagin. The inhibitory effect of Corilagin was also confirmed in RAW264.7 cells in which Stat6 was overexpressed based on the GV367-Stat6-EGFP lentiviral vector system and in which Stat6 was knock-downed by gene specific siRNAs. As a result, Corilagin prevented increases in the protein level of Phospho-Stat6 (P-Stat6). Both the mRNA and protein levels of the downstream mediators SOCS1, KLF4, and PPARγ/δ were markedly suppressed after Corilagin treatment. Expression of ARG1 and FIZZ1/Retnla, Ym1, TGF-β and PDGF in serum were also inhibited by Corilagin. The pathological changes, area of granulomas of liver sections, and degree of hepatic fibrosis were significantly alleviated in the Corilagin group. The areas of CD68- and CD206-positive cells stained by immunofluorescence were significantly decreased by Corilagin. In conclusion, Corilagin can suppress post-parasiticide schistosome egg-induced hepatic fibrosis by inhibiting the Stat6 signalling pathway and provide a new therapeutic strategy for schistosomiasis liver fibrosis.

## Introduction

Schistosomiasis is a prevalent parasitic disease that afflicts more than 240 million people from 78 countries and territories (1). Schistosomiasis japonica is a major intravascular infection that has serious public health consequences in China (2). Chronic infection with Schistosoma japonicum can cause inflammation and severe fibrosis, leading to disease-related morbidity and mortality (3, 4). Over the past few decades, the anthelmintic drug Praziquantel has been the basis of control and treatment for all forms of schistosomiasis (5–7). However, Praziquantel is quite ineffective against the young immature schistosome (8, 9). Schistosoma eggs are deposited in the liver of human hosts and give rise to inflammatory granulomatous in the chronic infection phase with liver fibrosis (10–13). Researchers found that during Schistosoma infection, macrophages respond to Th2 cytokines, and consequently, both IL-4 and IL-13 are abundant in chronic infection and are associated with liver fibrosis (14, 15). These macrophages are now commonly termed as “alternatively activated macrophages” (AAMϕ) or M2 macrophages (16). IL-4 and IL-13 signals mediate the M2 macrophage phenotype by activating Stat6 (17–19). Therefore, to find additional treatments for Schistosomiasis, targeting alternatively activated macrophages may be a strategy for treating this infection.

Corilagin (beta-1-O-galloyl-3, 6-(R)-hexahydroxydiphenoyl-D-glucose) is a tannin with the molecular formula C_27_H_22_O_18_ and is isolated from a variety of plants (20, 21). Corilagin has been shown to have an extensive pharmacological efficacy. A study demonstrated that *Phyllanthus urinaria L* containing Corilagin had antiviral activity that was marked by reduced cytotoxicity induced by EV71 or CA16 (22). The antinociceptive and anti-inflammatory effects of Corilagin, which is isolated from *Geranium bellum*, make this compound a candidate for the treatment of mild pain (23). The activity of Corilagin as an anti-hyperalgesic was verified in an experiment based on mouse models (24). Moreover, several studies have been carried out to obtain experimental evidence that shows that Corilagin has certain effects on multi-target and multi-channel treatments of hepatocellular carcinoma and ovarian cancer (25–27). It is interesting to note that it was confirmed that Corilagin attenuates lung cell damage induced by cigarette smoke (28). Some of our previous studies on Corilagin have confirmed its effects on ameliorating sepsis (21) as well as its anti-oxidative and anti-inflammatory effects (29) and suppressive effects on HSV-1 encephalitis (30). Moreover, our studies have demonstrated that IL-13 is associated with the multi-target effects of Corilagin on relieving schistosome egg-induced liver fibrosis (12, 32, 33). On the basis of the above findings, further study of Corilagin is warranted.

Thereby, the aim of this research is to explore the mechanism of Corilagin-associated inhibition of post-parasiticide schistosome egg-induced liver fibrosis through the Stat6 signalling pathways in hepatic alternative activated macrophages *in vitro* and *in vivo*.

## Materials and methods

### Chemicals and Reagents

The Corilagin Standard substance (purity>99%) for cell experiments was provided by China National Institute for the Control of Pharmaceutical and Biological Products (Beijing, China). Corilagin for animal experiments (purity≥80%) was obtained from Chengdu Purechem-standard biotechnology limited company (Chengdu, China). RAW264.7 cells were purchased from the China Center for Type Culture Collection (CCTCC) (Wuhan, China). DMEM medium was purchased from Gibco (Grand Island, NY, USA). Fetal bovine serum (FBS) was purchased from Zhejiang Tianhang Biological Technology Co., Ltd (Hangzhou, China). IL-13 and IL-4 were purchased from Peprotech (Rocky Hill, NJ, USA). The cell counting kit-8 (CCK-8) was purchased from Dojindo (Kumamoto, Japan). The rabbit anti-mouse KLF4 and PPARγ antibodies were purchased from Santa Cruz Biotechnology (Santa Cruz, CA). The rabbit anti-mouse SOCS1 antibody was purchased from ABclonal Biotech Co., Ltd. (Wuhan, China). The rabbit anti-mouse Stat6, PPARδ, CD68 and CD206 antibodies were purchased from Proteintech Group, Inc. (Chicago, USA). The rabbit anti-mouse Phospho-P65 (P-P65) antibody was purchased from Cell Signaling Technology (CST) (Boston, MA, USA). The rabbit anti-mouse Phospho-Stat6 (P-Stat6) antibody was purchased from Abcam PLC (Cambridge, UK). Horseradish peroxidase (HRP)-labelled goat anti-rabbit IgG was provided by Boster Immunoleader (Fremont, CA). The RNAiso and real-time RT-PCR kits were purchased from Takara (Dalian, China). The Mouse ARG1, Fizz1, Ym1, TGFβ and PDGF ELISA kits were purchased from Elabscience Biotechnology Co., Ltd (Wuhan, China). All primers were synthesized by Tsingke Biological Technology (Wuhan, China). Oncomelania hupensis was provided by Nanjing Center for Disease Control and Prevention (Nanjing, China).

### Cell culture

DMEM medium mixed with 10% foetal bovine serum was used to culture RAW264.7 in an incubator at 37°C, 5% CO_2_ and saturated humidity. Every 2–3 days, the supernatant was changed or the cells were passaged. Then, the normal control group, model group, Corilagin (100 mg/L, 50 mg/L and 25 mg/L) groups, Praziquantel (PZQ) group and Levofloxacin (LVFX) group were established. After the cells were passaged in 6-well plates or 96-well plates (CCK8 experiments and preliminary lentiviral transfection experiments) and cultured to 70% density, excluding the normal group, they were treated with IL-13 and IL-4 cytokines. After 24 h, Corilagin was diluted in DMEM medium to concentrations of 100 mg/L, 50 mg/ L and 25 mg/ L to interfere with the IL-13/IL-4 cellular model. After 24 h, the supernatants and cells were harvested for ELISA, quantitative real-time PCR and western blotting. Cells stimulated with IL-13 and IL-4 without any intervention were observed as a model group, whereas cells incubated in DMEM medium alone served as the normal control.

### IL-13/IL-4 administration in RAW264.7 for cell model establishment

Interleukin (IL)-13 and interleukin (IL)-4 were used to generate M2 macrophages to establish the cellular-model. The optimal dosage and timing of IL-13 and IL-4 for activating RAW264.7 were tested. After cells were passaged in 6-well plates and cultured to 70% density, the cell supernatants were removed and IL-13 (25ng/ml) and IL-4 (25ng/ml) were added to the wells. After 0 h, 6 h, 12 h, 24 h, 36 h, or 48 h, cells were harvested to detect the level of the downstream signalling molecules KLF4 and SOCS1 by quantitative real-time PCR and western blotting to explore the timing of their activity. A cytokine dose experiment was performed. After passaging cells in 6-well plates and culturing them to 70% density, the cell supernatants were removed and IL-13 and IL-4 at concentrations of 0 ng/ml, 25 ng/ml, 50 ng/ml, and 100 ng/ml were added to the wells. After 24 h, cells were harvested to detect the level of the downstream signalling molecules KLF4 and SOCS1 by quantitative real-time PCR and western blotting to determine the optimal dosage of IL-13 and IL-4 (31, 34).

### CCK8 Assay for Measuring the Cytotoxicity of Corilagin on RAW264.7 Cells

Cells were inoculated in 96-well (1×10^4^) plates. Then, Corilagin was added at different concentrations (0μg/ml, 25μg/ml, 50μg/ml, 100μg/ml, 200μg/ml, 300μg/ml, 400μg/ml, 500μg/ml). After 24 h, 10 μl of CCK8 was added to each well, followed by a 4-h incubation at 37°C before abeyance. The viability of the cells was measured at 490 nm using a microplate reader according to the manufacturer’s instructions.

### Knockdown of Stat6

Mouse Stat6 siRNA (sense 5’-CCTGCAACCATCTCCTTAT-3’) was synthesized by RiboBio Co., Ltd. GenePharma Co. Ltd (Guangzhou, China). RAW264.7 cells were placed onto 6-well plates and transfected with siRNA duplexes using Lipofectamine 2000 (Invivogen, San Diego, USA) according to the manufacturer’s instructions. The medium was replaced 6 h later, and then, cells were grown for up to 48 or 72 h. At 24 h before harvest, the intervened cells were treated with Corilagin, Praziquantel (PZQ) or Levofloxacin (LVFX). Knockdown of Stat6 was confirmed by real-time PCR analysis

### Over-expression of Stat6

For overexpression, Stat6 lentiviral vectors were synthesized by GeneChem Co, Ltd. (Shanghai, China). GV367-Stat6/NC-EGFP were transfected into the 293T Cell line, and the viral supernatant was harvested after 48 h. RAW264.7 cells were seeded onto 6-well plates and transfected with lentivirus with a multiplicity of infection (MOI) of 80 according to the manufacturer’s instructions. The medium was replaced 6 h later, and then, the cells were grown for up to 72 h. Then, the lentivirus intervened cells were treated with Corilagin, PZQ or LVXF for another 24 h. Overexpression of Stat6 was confirmed by real-time PCR analysis (30).

### Animals

Forty-two six-week-old specific pathogen-free (SPF) Balb/c male mice (18-22 g) were purchased from Hubei Provincial Center for Disease Control and Prevention (Wuhan, China). Mice were maintained under standard laboratory conditions, and animals were fed a normal diet and water ad libitum as previously described (32). All experiments and animal care abided by internationally accepted principles and the Guidelines for the Care and Use of Laboratory Animals of Huazhong University of Science and Technology and were approved by the Ethics Committee of Union Hospital, Tongji Medical College, Huazhong University of Science and Technology (IACUC Number: S759)

### Animal grouping and treatment

All mice were randomly assigned to the following treatment groups with six mice per group: the normal group, model group, Corilagin low dosage group (20 mg/kg/d), Corilagin medium dosage group (40 mg/kg/d), Corilagin high dosage group (80 mg/kg/d), Praziquantel (PZQ) group and Levofloxacin (LVFX) group. All of the groups except the normal group were infected with 25±5 cercariae of Schistosoma japonicum through the abdominal skin as described in our previous research (35). After four weeks, the infected groups were intragastric administered Praziquantel (PZQ) (500 mg/kg/d) for one week to kill adult worms. Then, the infected groups received three weeks of the corresponding medications. Mice in the low, middle, and high-dose Corilagin groups were administered 20, 40, and 80 mg/kg Corilagin (concentration of 2, 4, 8 mg/ml, the feeding volume was 10 ml/kg) by gavage, and mice in the PZQ group (500 mg/kg/d) and LVXF group (100 mg/kg/d) also received three weeks of treatment. Normal saline was administered to the model and normal groups with the same volume of 10 ml/kg over the three-week treatment period. All mice were sacrificed at four weeks after administration.

### Specimen collection and liver histological studies

The protocol for specimen collection was previously described (33). Following induction of anaesthesia with 6% chloral hydrate via intraperitoneal injection, the mouse abdomen was opened and the abdominal aorta was separated. Then, 2 ml of ocular blood was collected in a coagulant test tube. Serum was obtained after centrifugation at 3500 rpm/min for 15 min and stored at −20°C until testing. Subsequently, the livers were collected with an aseptic, RNase-free apparatus. After washing them with normal saline, tissues were stored in liquid nitrogen.

### ELISA for measuring ARG1 in the supernatant and PDGF, Fizz1, Ym1, TGFβ in serum

The procedures for measuring ARG1 in the supernatant and PDGF, Fizz1, Ym1, TGFβ in serum were carried out according to our previous study (36, 37). Expression of ARG1 in the supernatant and PDGF, Fizz1, Ym1, and TGFβ in serum was determined by sandwich ELISA. The supernatant and serum were collected and assayed for ARG1, Fizz1, Ym1, PDGF and TGFβ by the respective ELISA kits according to the manufacturer’s instructions.

### Quantitative real-time PCR for detecting mRNA expression

The procedure for quantitative real-time PCR was described in our previous study (33). Total RNA in liver tissues or cultured cells was isolated using RNAiso Plus according to the manufacturer’s protocol. The cDNAs were produced with the PrimeScriptTM RT reagent Kit and were incubated at 37 °C for 15 min and 85°C for 5 s. Real-time PCR reactions were performed in a Step One Plus device at 95 °C for 10 s followed by 40 cycles of 95 °C for 5 s and 60°C for 20 s according to the instructions for the SYBR Premix Ex Taq kit. The 2^−ΔΔCt CT^ method was performed to analyse the results (38). The primer sequences (5′→3′) were as follows.

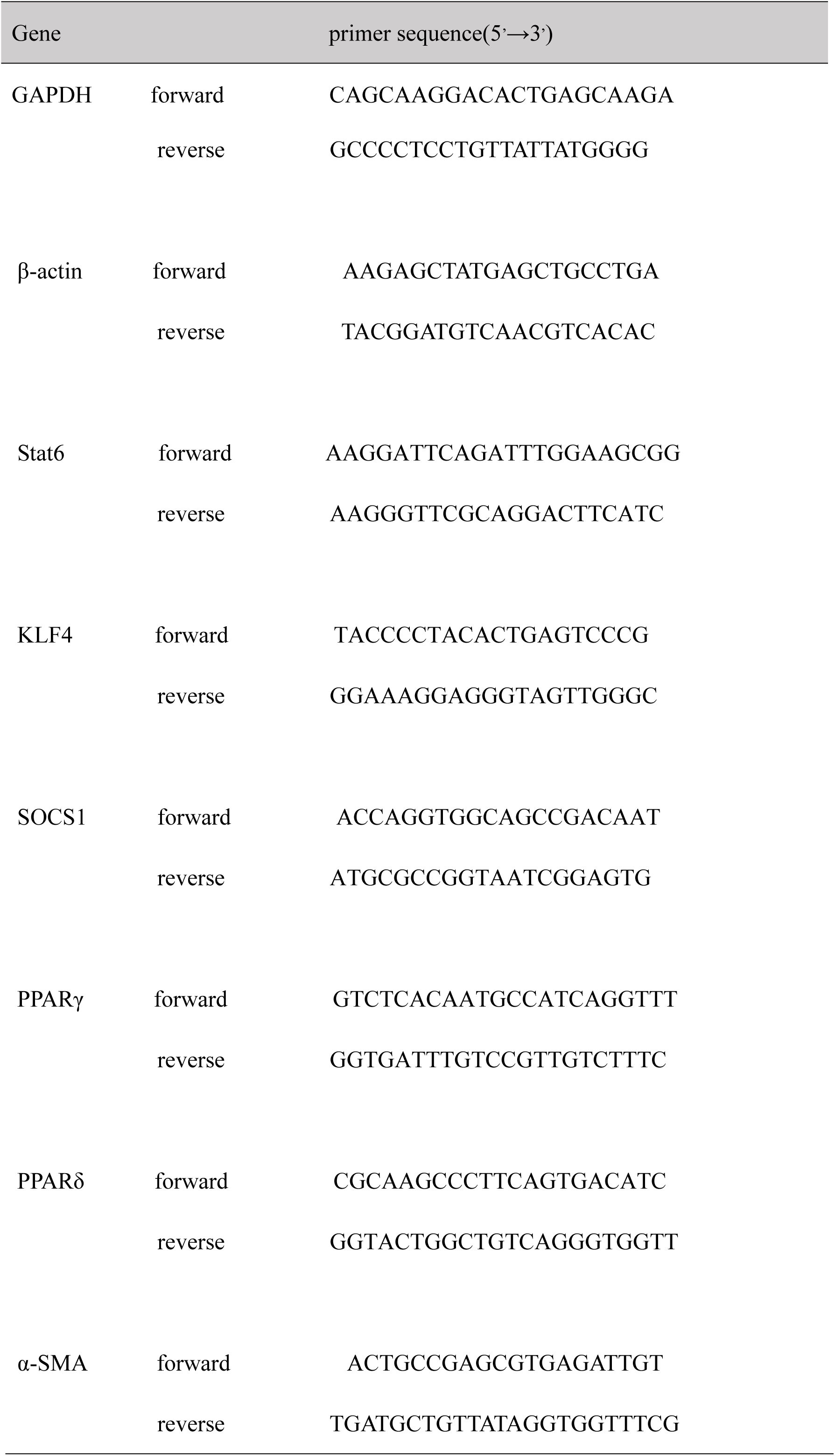

### Western blotting for measuring protein expression

The western blotting protocol followed our previous study (39). Proteins extracted from either liver tissue or cultured RAW264.7 cells were analysed by western blotting. The protein concentration was determined using the Bicinchoninic acid (BCA) method. To each tube, an equivalent volume of 2× sodium dodecyl sulphate (SDS) loading buffer (100 mM Tris–HCl, pH 6.8, 4% SDS, 20% glycerine, 10% beta-mercaptoethanol, and 0.2% bromophenol blue) was added and mixed. The mixtures were then denatured at 95 °C for 10 min. Then, the proteins were separated on SDS-PAGE gels and transferred onto a nitrocellulose filter membrane. The membranes were blocked overnight with 5% non-fat milk in phosphate-buffered saline (PBS) and probed with the indicated antibody before being washed three times in Tris-buffered saline with Tween (TBST), followed by incubation with an HRP-labelled secondary antibody. After the membranes were further washed with TBST, electrochemiluminescence (ECL) was used to identify the immunoreactive bands. Densitometry analysis of the immunoreactive bands was performed using the Fuji ultrasonic-Doppler velocity profile (UVP) system and the Image J program.

### Pathological examination of liver tissue

The procedure of H&E staining and Masson staining was conducted as reported in our past experiment (40). Liver tissues were fixed in 4% formalin and embedded in paraffin. Four-micrometre serial sections were obtained for H&E staining and Masson staining to assay pathological changes in the liver to compare the area of egg granulomas. Measurement of granulomas in the livers was defined as the average area.

### Immunofluorescence (IF)

The slides were washed three times with PBS and were incubated with 10% normal goat serum for 20 min at 37°C. Then, the slides were incubated at 4°C overnight with a primary antibody (CD68 1:100, CD206 1:100). The slides were washed with PBS and exposed to an Alexa Fluor fluorochromes-conjugated goat IgG secondary antibody (Peprotech, Rocky Hill, NJ, USA) and then diluted at 1:200 at 37 °C for half an hour without light. The slides were washed in PBS three times and were then incubated with DAPI for 10 min and mounted (41, 42).

### Statistical analysis

Data are presented as the mean ± standard deviation (SD). Data were compared between groups by ANOVA and the SNK test. Statistical significance was considered when P<0.05 or P <0.01 (43). Statistical analyses were conducted with SPSS 12.0 software.

## Results

### Cytokine treatment and cytotoxicity of Corilagin

As shown in Fig1, the levels of KLF4 and SOCS1 in RAW264.7 cells reached their maximum at 24 h after IL-4 (25ng/ml) stimulation. For the IL-13-induced group (25ng/ml), the levels of KLF4 and SOCS1 reached their maximum at 36 h after stimulation, and expression at 24 h also reached high levels and showed no significant difference relative to the levels after 36 h of treatment. Thus, the optimal timing both for IL-13 and IL-4 to activate RAW 264.7 macrophages was 24 h. Moreover, there was a remarkable increase in the expression of KLF4 and SOCS1 at 25 ng/ml and 100 ng/ml after IL-13 treatment and at 50 ng/ml and 100 ng/ml after IL-4 stimulation. Therefore, concentrations of 25 ng/ml for IL-13 activation and 50 ng/ml for IL-4 treatment were chosen as the optimal dosages for subsequent experiments. As shown in Fig. 2, the concentration at which Corilagin inhibited RAW264.7 cell growth by 50% (IC50) was approximately 160 μg /ml for 24 h. The viability of cells approached 80% at 100 μg/ml Corilagin after 24 h. Therefore, we used Corilagin at 100 μg/ml, 50 μg/ml, and 25 μg/ml to treat RAW 264.7 cells for 24 h.

**Figure 1.**
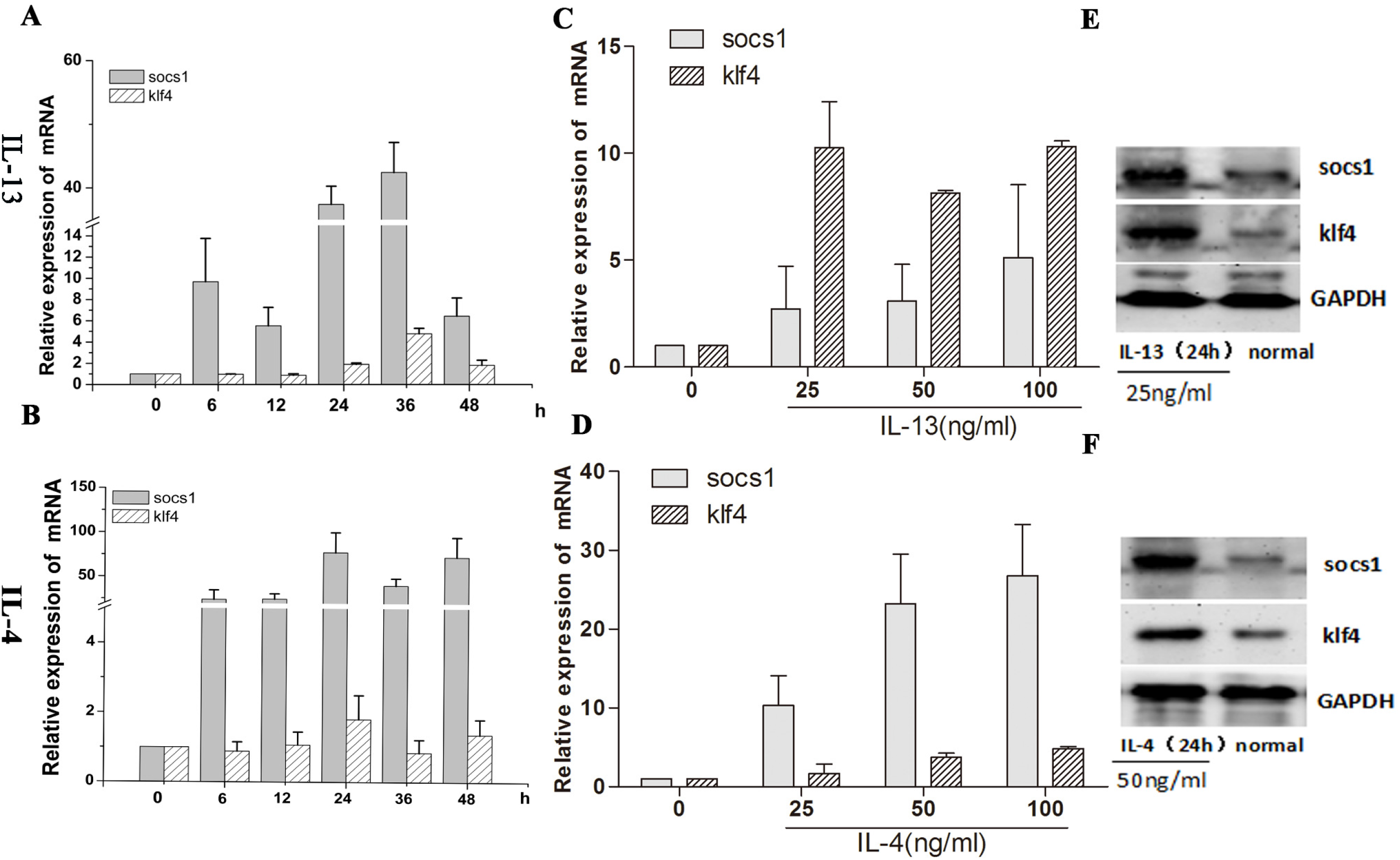
The level of SOCS1, KLF4 in IL-4/IL-13-induced RAW264.7 cells based on time and dosage of IL-13 and IL-4. **(A-B)** The mRNA level was detected by real-time PCR at 0, 6, 12, 24, 36, and 48 h after IL-4/IL-13(25ng/ml) treatment. **(C-D)** After the timing was determined, the effects of IL-13/IL-4 on SOCS1 and KLF4 at concentrations of 25, 50, 100 ng/ml were detected by real-time quantitative PCR after 24 h of treatment. **(E-F)** The effects of IL-13 (25 ng/ml, 24 h) and IL-4 (50 ng/ml, 24 h) on SOCS1 and KLF4 were confirmed by western blotting. Data shown are the mean ± SD from 3 independent experiments.

**Figure 2.**
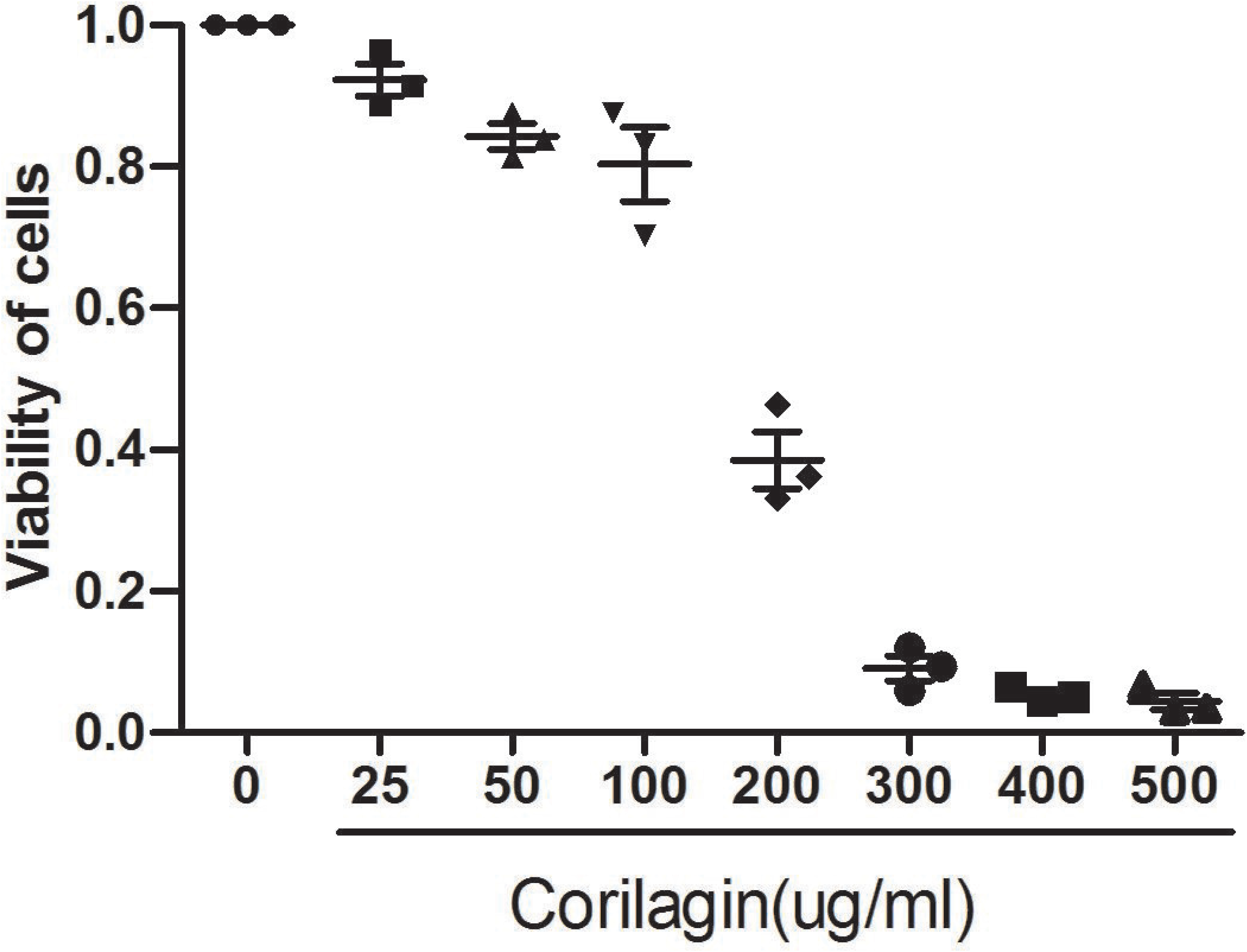
Cytotoxicity of Corilagin on RAW264.7 cells. The cytotoxic effect of Corilagin was evaluated by the CCK8 assay. Data shown are the mean ± SD from 3 independent experiments.

### Evaluation of ARG1 expression in the supernatant by ELISA

M2 macrophages express increased levels of arginase-1 (Arg1). Thus, Arg1 is associated with M2 macrophages or M2-like cells and was detected as a definitive M2 marker (44). In Fig. 3, after IL-4/IL-13 stimulation, mRNA expression of Arg1 was much higher than that in the normal group and M2 macrophages were activated. When the Corilagin treatment groups were compared with the model group, the level of ARG1 was significantly lower (P<0.05) and was dose-dependent (P<0.05).

**Figure 3.**
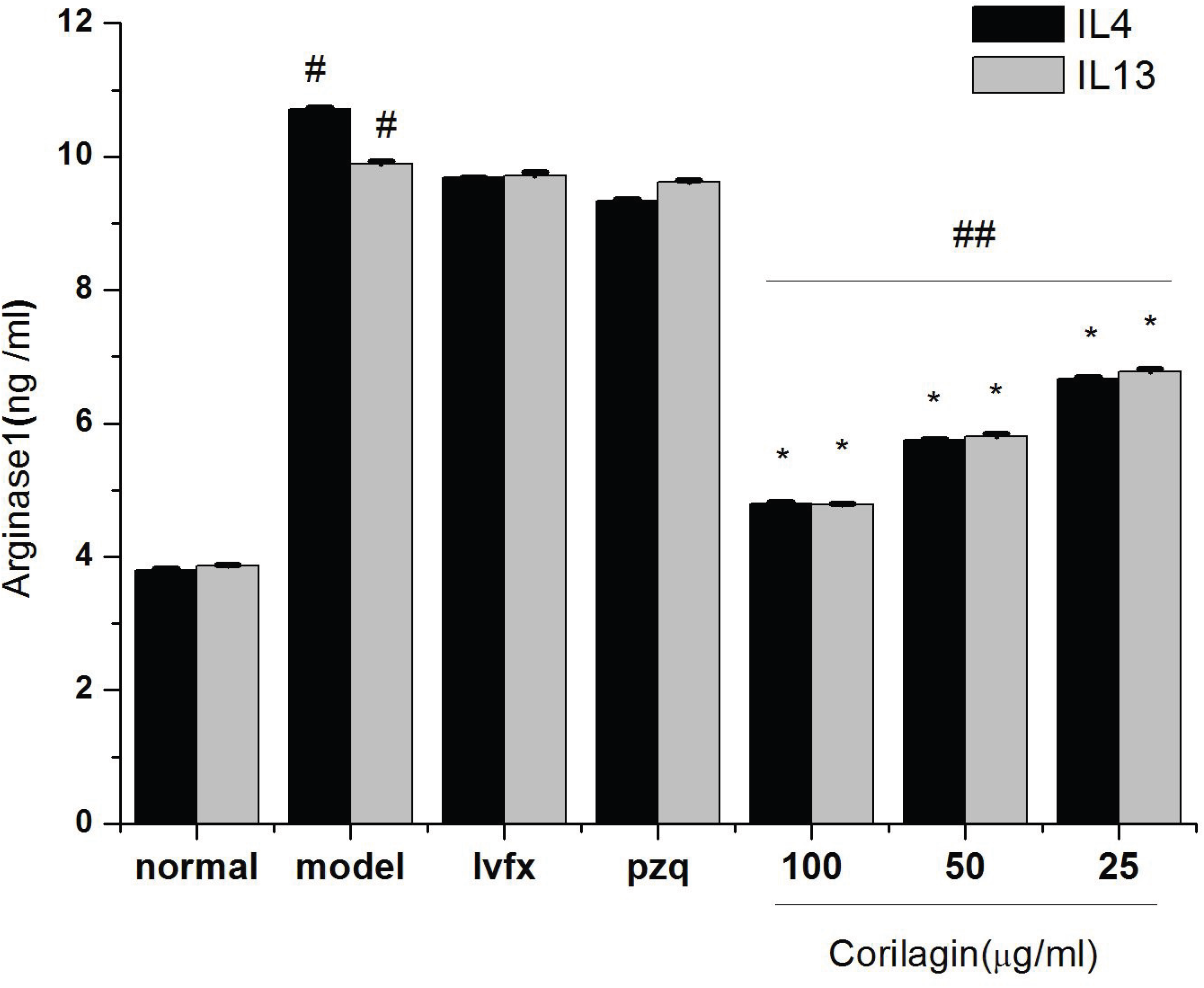
Effect of Corilagin on ARG1 in supernatant. Effect of Corilagin on the expression of ARG1 according to ELISA. Data shown are the mean ± SD from 3 independent experiments. ^#^P< 0.05 *vs*. normal group; *P< 0.05 VS model group; as determined by Student’s t-test; ^##^P< 0.01 determined by One-way ANOVA.

### Effect of Corilagin on Stat6 and downstream molecules in RAW 264.7 cells after IL-13 /IL-4 stimulation

As shown in Fig. 4, IL-13 and IL-4 remarkably elevated the protein expression of P-Stat6 compared with the normal group. However, there was a significant decrease in the protein expression of P-Stat6 after Corilagin treatment. Meanwhile, IL-13/IL-14 and Corilagin had the opposite effect on the protein expression of P-P65. Moreover, the mRNA and protein levels of the downstream molecules KLF4, SOCS1, PPARγ and PPARδ were also significantly increased in cells stimulated by IL-13 and IL-4 (P <0.05). However, Corilagin had a remarkable inhibitory effect on the expression of KLF4, SOCS1, PPARγ and PPARδ (P <0.01 or 0.05) that was concentration-dependent (P <0.05).

**Figure 4.**
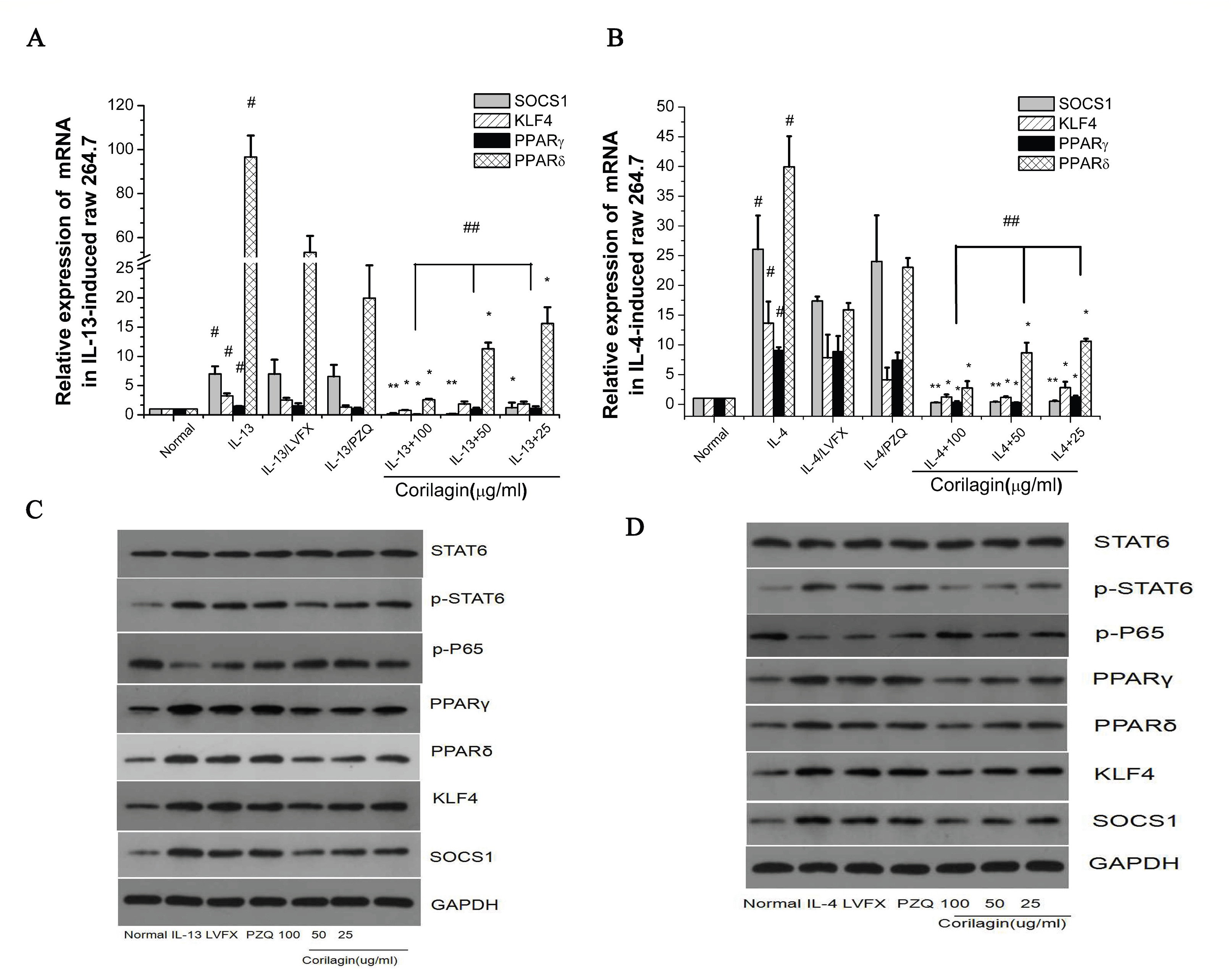
Effect of Corilagin on Stat6 and downstream molecules in IL-13-induced/IL-4-induced RAW264.7 cells. **(A-B)**The mRNA levels of downstream molecules, including PPARγ, PPARδ, KLF4 or SOCS1, were detected by real-time quantitative PCR. **(C-D)** The protein levels of P-Stat6, P-P65, SOCS1, KLF4, PPARδ, and PPARγ were measured by western blot. Data shown are the mean ± SD from 3 independent experiments. ^#^P< 0.05 VS normal group, *P< 0.05, ** P< 0.01 *VS* IL-13-induced group/IL-4-induced group; As determined by Student’s t-test; ^##^P< 0.05 determined by One-way ANOVA.

### Effect of Corilagin on downstream signalling molecules in RAW 264.7 cells after Stat6 overexpression

We generated lentivirus-mediated Stat6 overexpressing RAW264.7 cells, and expression of Stat6/P-Stat6 was confirmed after 72 h (Figure 5B and Figure 5D). Green Fluorescent Protein (GFP) was observed with a fluorescence microscope after lentivirus was introduced to cells for 36 h and 72 h (Figure 5A). Compared with the normal group, the expression levels of KLF4, SOCS1, PPARγ and PPARδ were significantly increased in the Stat6 overexpression group (P <0.01 or 0.05) (Figure 5C-5D). Twenty-four hours after IL-13/IL-4 intervention, cells were treated with Corilagin (25, 50 or 100 μg/ml) for 24 h. As shown in Fig.6, expression of KLF4, SOCS1, PPARγ and PPARδ was reduced in the Corilagin-treated group compared with that in the IL-13/IL-4-induced group (P <0.01 or 0.05). Therefore, we found that Corilagin could inhibit downstream molecules in RAW264.7 cells when Stat6 was overexpressed.

**Figure 5.**
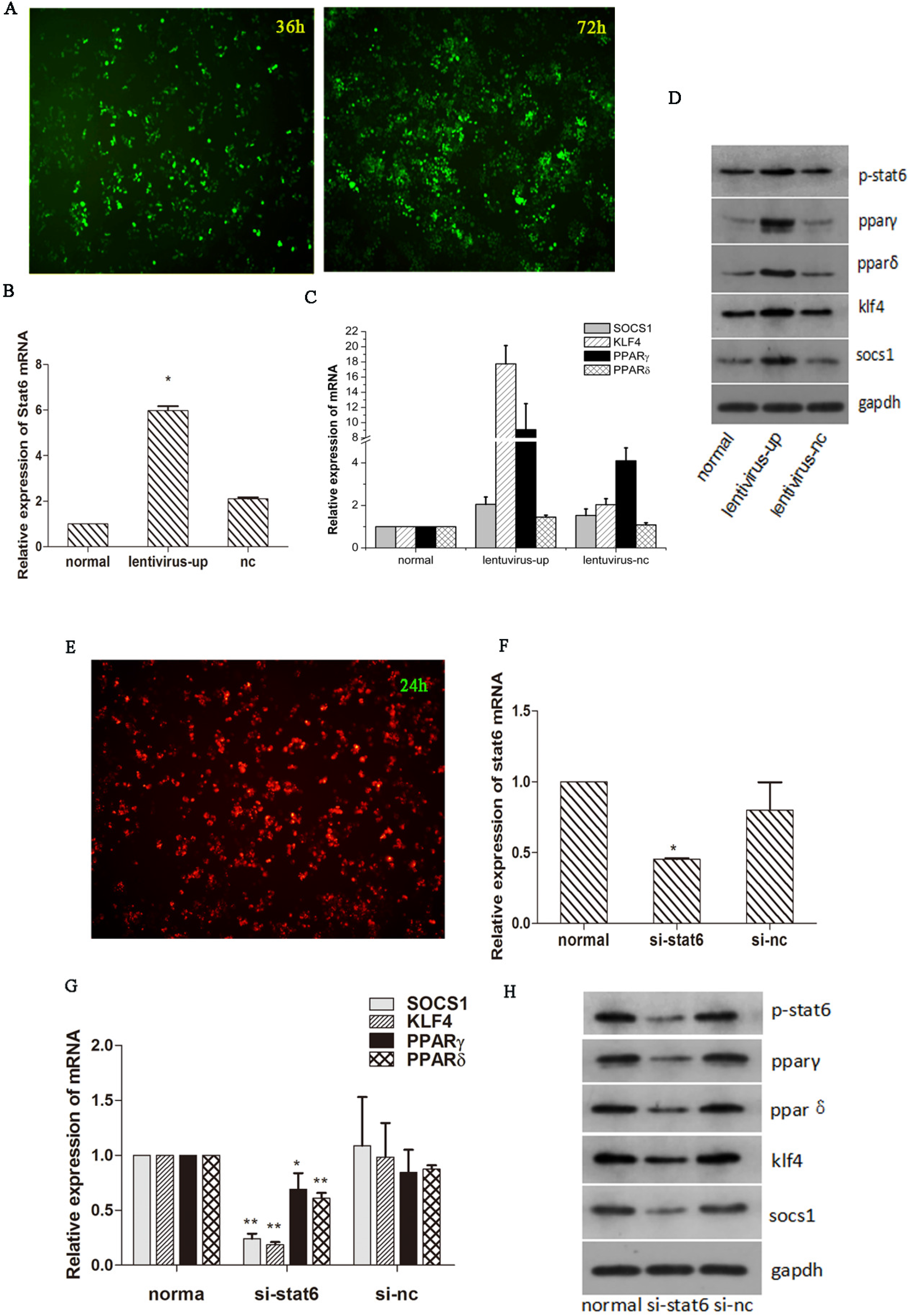
Up-regulatio/Down-regulation of Stat6 in RAW264.7 cells via the lentiviral vector/. **(A)** Expression of GFP was observed with fluorescence microscope after lentivirus was introduced into cells for 36 h and 72 h. **(B-C)** The level of Stat6 and downstream molecules after transfection was confirmed by real-time quantitative PCR. **(D)** Expression of protein was assayed by western blotting. **(E)** Expression of RFP was observed with a fluorescence microscope after siRNA was introduced into cells for 24 h. **(F-G)** The levels of Stat6 and downstream molecules after transfection were confirmed by real-time quantitative PCR. **(H)** Expression of protein was assayed by western blotting. Data shown are the mean ± SD from 3 independent experiments. *P< 0.05, ** P< 0.01 *VS* normal group; as determined by Student’s t-test;

### Effect of Corilagin on downstream signalling molecules in RAW 264.7 cells after Stat6 knocked-down

We used siRNA to silence the expression of Stat6 in RAW264.7 cells. The expression of Red Fluorescent Protein (RFP) was observed through a fluorescence microscope after siRNA was introduced into cells for 24 h (Figure.5E). The expression levels of Stat6/P-Stat6 as well as the downstream molecules KLF4, SOCS1, PPARγ and PPARδ are shown in Fig 5F-5H and underwent a significant decrease compared with levels in the group without Stat6 knockdown (P <0.01 or 0.05). We used IL-13 and IL-4 to treat transfected RAW264.7 cells for 24 h and then treated cells with Corilagin to observe its inhibitory impact on the downstream molecules of the Stat6 signalling pathways in Stat6 knocked-down cells. As shown in Fig. 6, the mRNA levels of KLF4, SOCS1, PPARγ, and PPARδ and protein levels of P-Stat6, KLF4, SOCS1, PPARγ, and PPARδ were significantly suppressed by Corilagin (P<0.01 or 0.05).

**Figure 6.**
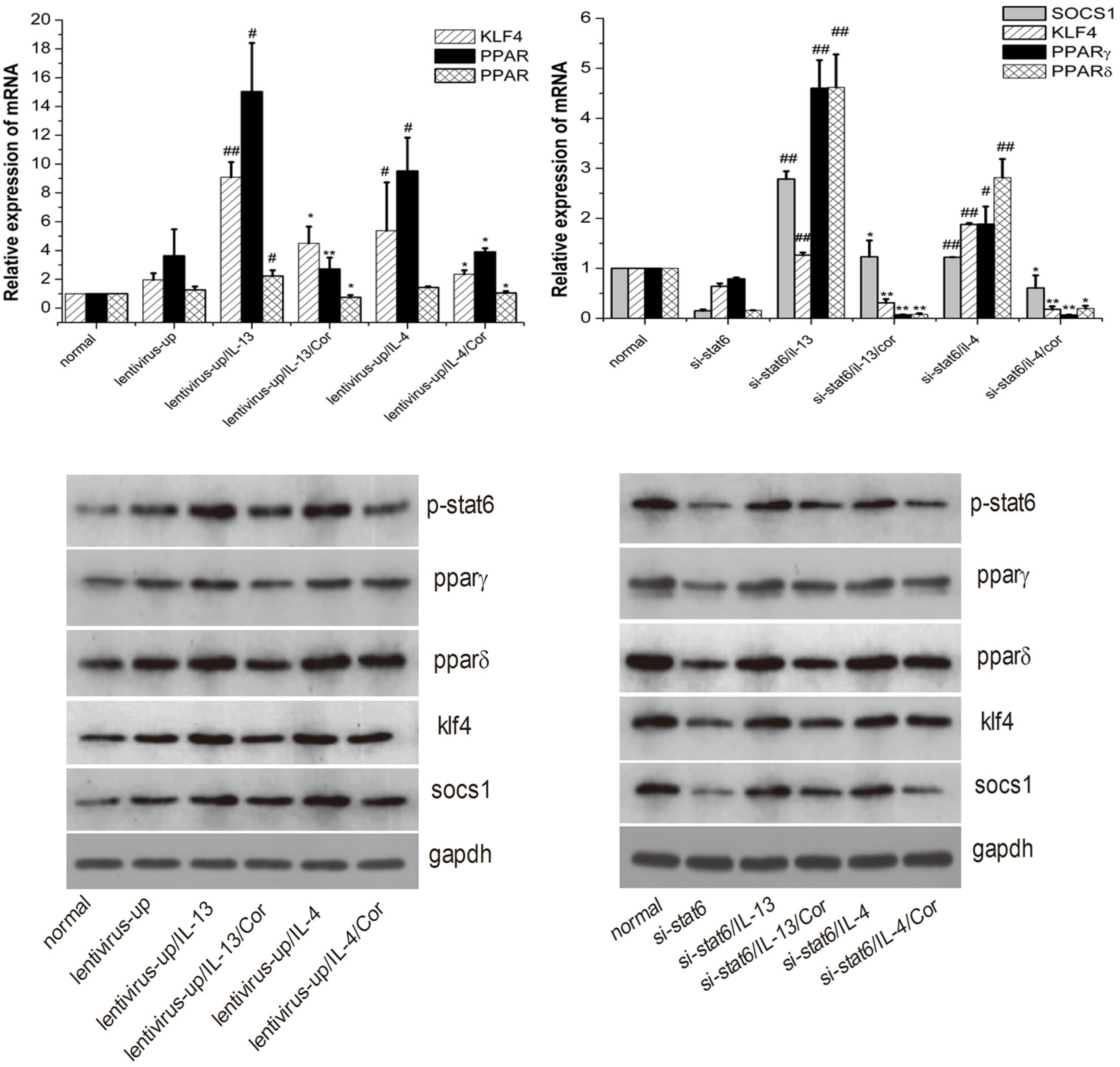
Expression of Corilagin on downstream signalling molecules in IL-13/IL-4-induced RAW 264.7 cells after Stat6 over-expression or knockdown. **(A/C)** The mRNA levels of downstream molecules, including SOCS1, KLF4, PPARδ, and PPARγ, were detected by real-time quantitative PCR. **(B/D)** The protein levels of P-Stat6, SOCS1, KLF4, PPARδ, and PPARγ were measured by western blot. Data shown are the mean ± SD from 3 independent experiments. ^#^P< 0.05, ^##^P < 0.01 VS transfected group, *P< 0.05, ** P< 0.01 *VS* IL-13/IL-4-induced group; as determined by Student’s t-test;

### Effects of Corilagin on M2 macrophages in the liver by IF

The M2 macrophage mannose receptor-1 (CD206) can effectively distinguish between M1 and M2 macrophage subtypes (45). Evidence indicates that CD68 is expressed in all macrophages, and it has been labelled as a histological marker of macrophage lineage cells (46). We used two indictors to analyse the distribution and expression of M2 macrophages in the liver with or without treatment of Corilagin (47). As shown in Fig.7, immunofluorescence of CD206^+^ and CD68^+^ in the liver of the model group showed localization in the granuloma and nodular areas, which demonstrated high expression of M2 macrophages in the liver of schistosome egg-induced hepatic fibrosis. After Corilagin interference, the level of CD206^+^ and CD68^+^ was significantly decreased. Meanwhile, there was little expression of CD206^+^ in the intact normal liver and cells. PZQ performed similarly to Corilagin at 20 mg/kg, and there were no distinct changes in the levels of CD206 and CD68 after LVFX treatment relative to the levels observed in the model group. Taken together, these results suggest that the alternatively activated (M2) macrophage phenotype was activated in the liver of hepatic fibrosis schistosome egg-induced model mice. Moreover, Corilagin is effective at inhibiting these M2 genes.

**Figure 7.**
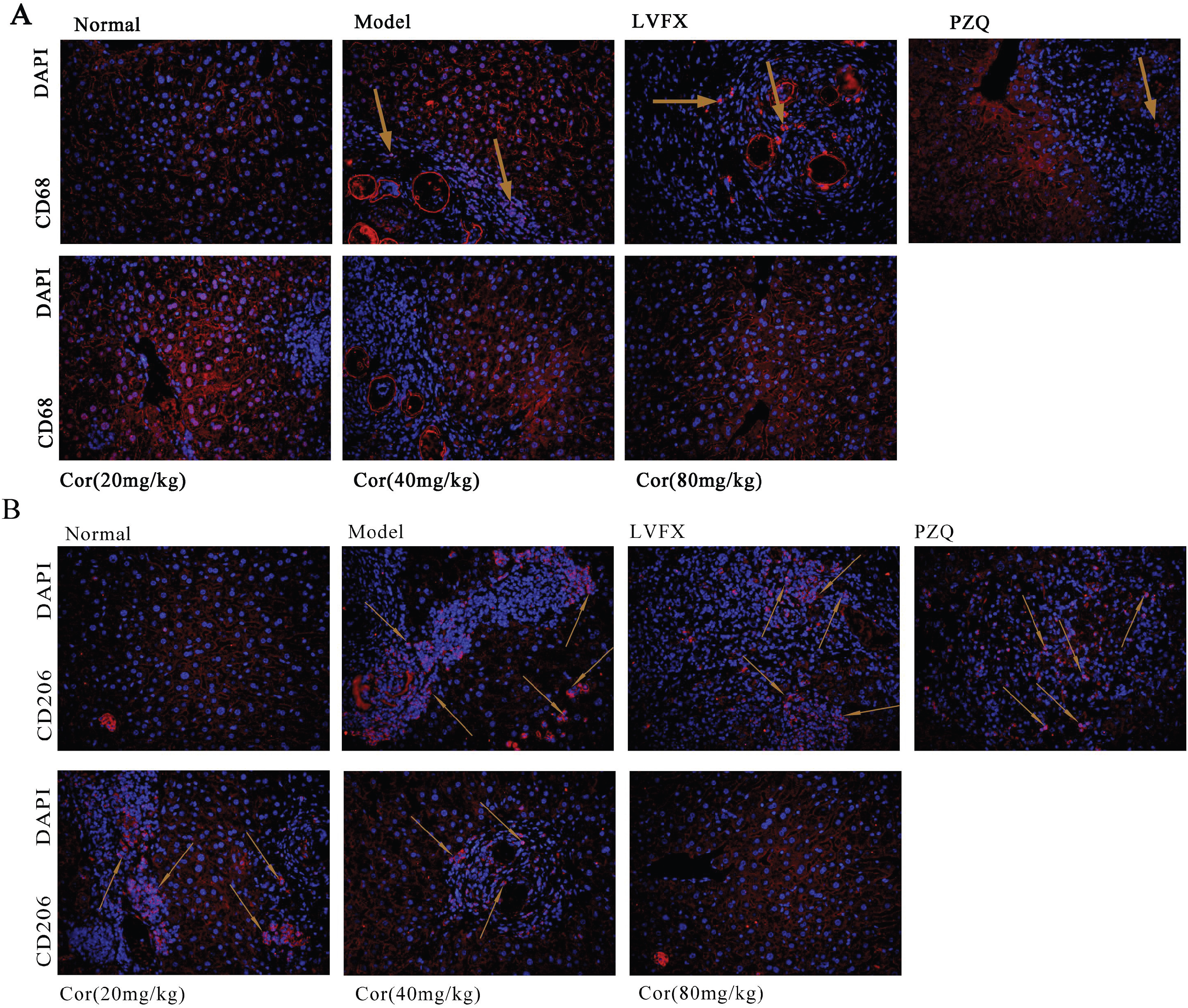
Immunofluorescence detection of CD206 and CD68 in liver tissue. **(A-B)** Localization and expression of mannose receptor CD68^+^ (red) and mannose receptor CD206^+^ (red). Nuclei were visualized by DAPI (blue) staining. Figures were captured at 400X magnification.

### Effects of Corilagin on Stat6 and downstream molecules in M2 macrophages in the livers of schistosomiasis-infected mice

As shown in Figure 8, compared with the normal group, the mRNA and protein levels of KLF4, SOCS1, PPARγ and PPARδ were significantly elevated in the model group (P<0.05). Meanwhile, there was an increase in the protein expression of P-Stat6, KLF4, SOCS1, PPARγ, and PPARδ as well as a decrease in the protein level of P-P65 in the model group. After Corilagin treatment, Stat6/P-Stat6, KLF4, SOCS1, PPARγ and PPARδ were significantly inhibited at the mRNA level and protein level (P <0.01 or 0.05). However, protein expression of P-P65 was remarkably elevated by Corilagin.

**Figure 8.**
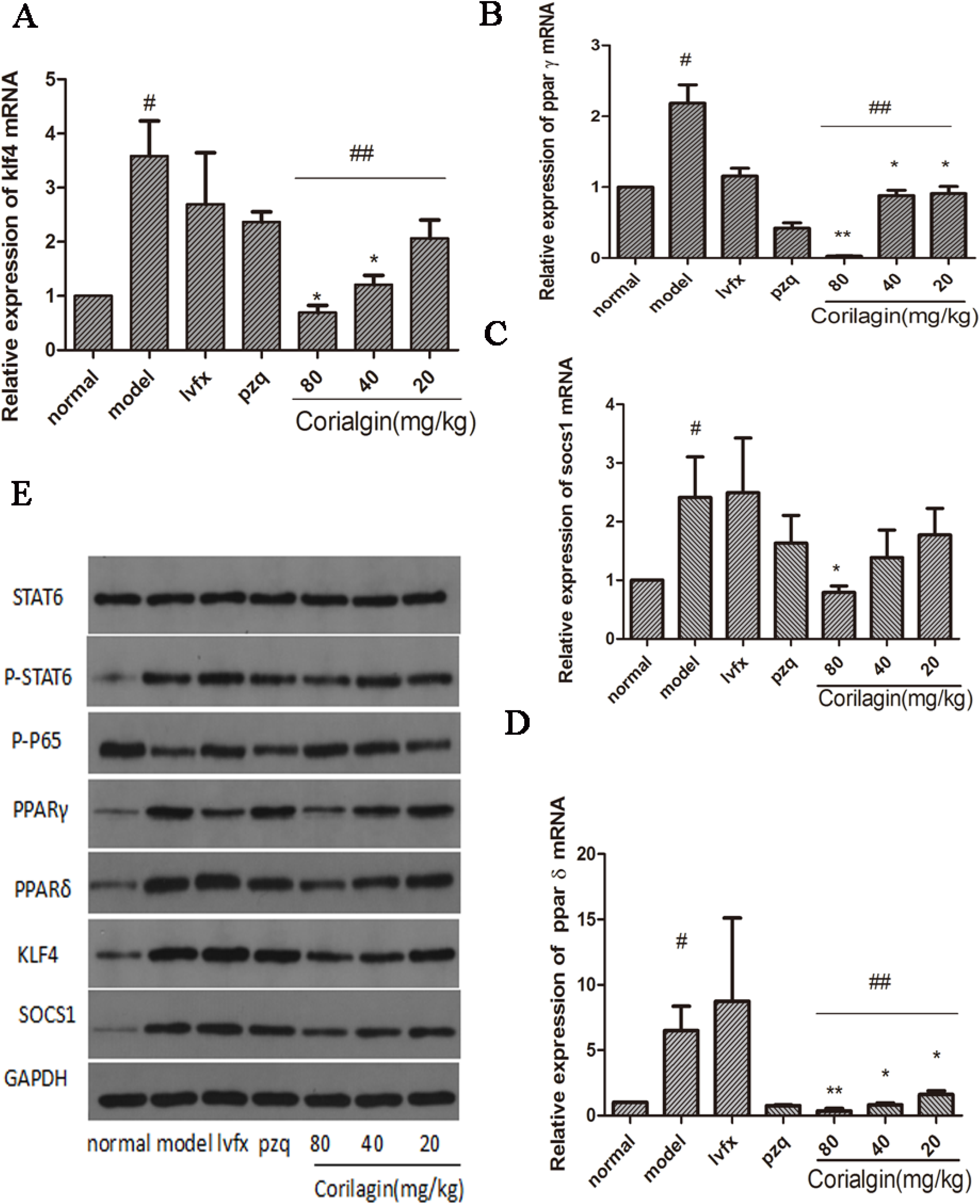
Effects of Corilagin on Stat6 and downstream molecules in M2 macrophages in the livers of schistosomiasis-infected mice. **(A-D)** The mRNA levels of downstream molecules, including SOCS1 (A), KLF4 (B), PPARδ (C), and PPARγ (D), were detected by real-time quantitative PCR. (E)The protein levels of P-Stat6, P-P65, SOCS1, KLF4, PPARδ, and PPARγ were measured by western blot. Data shown are the mean ± SD from 6 experiment mice. ^#^P< 0.05 *vs*. normal group, *P< 0.05, ** P< 0.01 *vs*. model group; as determined by Student’s t-test; ^##^P< 0.05 determined by One-way ANOVA.

### Effect of Corilagin on PDGF, Fizz1, Ym1 and TGFβ in liver tissue according to ELISA

M2 macrophages produce a number of factors, such as Fizz1, Ym1 and TGFβ, which are considered to be indicators of M2 macrophages (AAMϕ) (48, 49). PDGF is markedly overexpressed in fibrous tissues, and its activity increases with the degree of liver fibrosis (50). TGFβ is known as an inducer of fibrogenesis in hepatic fibrosis. As shown in Figure 9, the levels of the M2 macrophage genes Fizz1, Ym1, TGFβ and PDGF were significantly elevated in the model group compared with the levels in the normal group (P <0.01 or 0.05). Corilagin had a remarkable effect on inhibiting the Fizz1, Ym1, TGFβ and PDGF levels (P<0.05 or P<0.01). PZQ (40 mg/kg) had a similar effect as Corilagin on Ym1 and TGFβ. The LVFX group had no remarkable changes compared with the model group.

**Figure 9.**
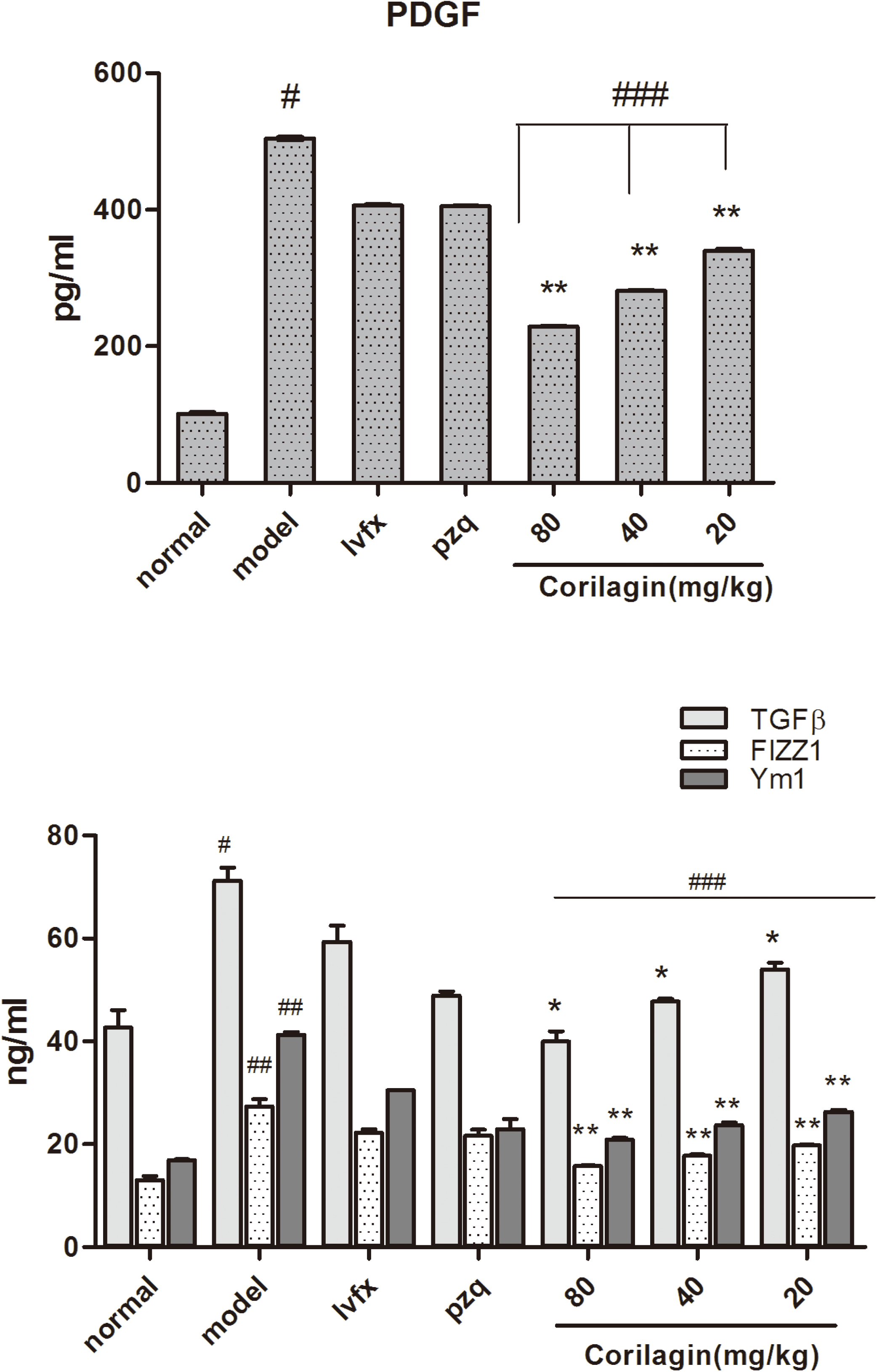
Effect of Corilagin on PDGF, Fizz1, Ym1, and TGFβ in the serum. Effect of Corilagin on expression of PDGF, Fizz1, Ym1, and TGFβ according to ELISA. Data shown are the mean ± SD from 6 experiment mice. ^#^P< 0.05, ^##^P< 0.01 *vs*. normal group; *P< 0.05, ** P< 0.01 *VS* model group; As determined by Student’s t-test; ^###^P< 0.01 determined by One-way ANOVA.

### Effect of Corilagin on pathological changes

As shown in Figure 10, in the normal group, hepatic tissue showed an intact network arrangement of hepatic lobules and cells; there was no egg granuloma or cellular fibrosis according to H&E and Masson staining. Compared with the normal group, the livers from the model group showed obvious pathological changes. Severe fibril aggregation and connective tissues surrounded large granulomas, and large necrosis foci were also observed in the liver, accompanied by some inflammatory cell infiltrates that were visible in both H&E and Masson staining. Corilagin eased and reduced inflammation, as indicated by a reduction in the average area of egg granulomas according to Masson staining (P<0.05 or P<0.01). (Figure 14C). By Masson staining, sections of liver tissue from chronically infected animals showed that egg granulomas accumulated in large numbers to form nodules, and these nodules were clustered, producing numerous variable-sized mass lesions in which the centre was necrotic and enclosed by fibrous cells. After treatment with Corilagin, the degree of inflammation was relieved and the mass lesions were remarkably reduced to different degrees. However, the effect of PZQ approached that between the Corilagin (20 mg/kg) group and Corilagin (40 mg/kg) group, and there was no significant effect after LVXF treatment. Fibrosis was scored as 5 grades according to the METAVIR scoring system (51, 52): F0, no fibrosis; F1, portal fibrosis without septa; F2, portal fibrosis with few septa; F3, numerous septa without cirrhosis; and F4, cirrhosis. The degree of hepatic fibrosis was detected to elevate the score F0(2^0^)=1, F1(2^1^)=2, F2(2^2^)=4, and F3(2^3^)=8 (12). The mRNA levels of Alpha α-SMA further demonstrated that the degree of hepatic fibrosis from the Corilagin group was significantly alleviated (P<0.05).

**Figure 10.**
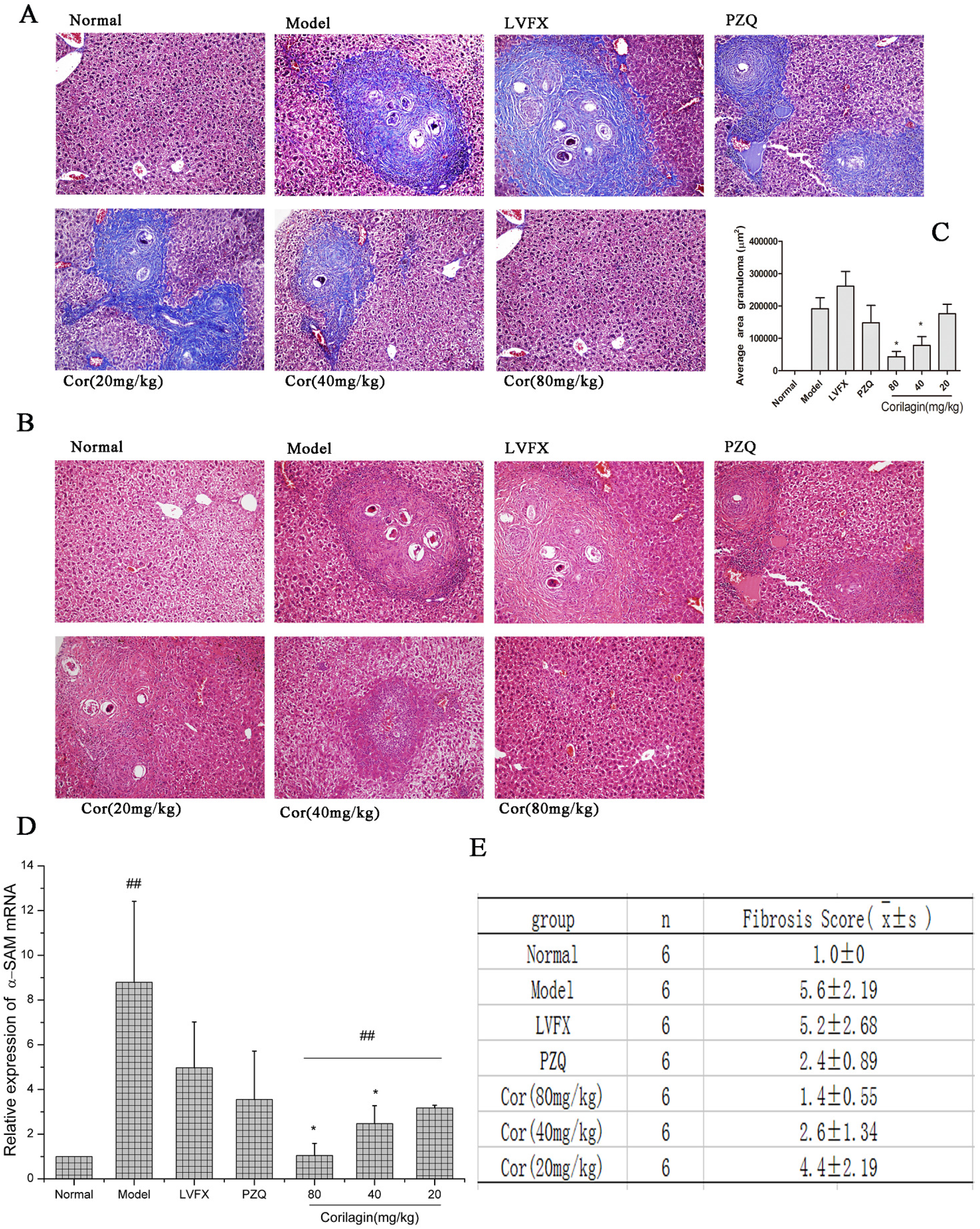
Effect of Corilagin on pathological changes. **(A-B)** Effect of Corilagin on pathological was examined by HE staining and Masson staining (×200). **(C)** Measurement of granulomas in the livers was defined as the average area. **(D)** The mRNA level of α-SMA was detected by real-time quantitative PCR. **(E)** liver fibrosis score. Data shown are the mean ± SD from 6 experiment mice. *P< 0.05 *vs*. model group; as determined by Student’s t-test; ^##^P< 0.05 determined by One-way ANOVA.

## Discussion

Liver fibrosis is a pathologic process that results from chronic liver diseases of various aetiologies, and it can lead to disease-related morbidity and mortality (3, 53). Chronic Schistosoma infection is one of the known aetiologies leading to liver fibrosis. Schistosomiasis is a major neglected tropical infectious disease that has serious public health consequences (54). In China, schistosomiasis japonica is a major health risk for more than 50 million people (55). Schistosomiasis morbidity is due to chronic infection with Schistosoma japonicum, which is characterized by liver fibrosis at chronic and advanced stages (56, 57). Schistosome egg-induced hepatic liver fibrosis results from chronic infection mainly due to the granulomatous inflammatory response against Schistosoma eggs. Praziquantel is the only drug that is effective against all Schistosoma species, including S. japonicum, and it is effective in acting against adult schistosome worms, but is ineffective in acting against immature worms and worm eggs (58). Schistosoma eggs reach the liver via the portal venous system and then they can cause an inflammatory reaction, leading to fibrosis (59). Therefore, finding a therapeutic target for schistosome egg-induced liver fibrosis may help lead to a more effective curative effect by impairing parasite eggs.

Studies found that alternatively activated macrophages (AAMϕ) or M2 macrophages compose 20% to 30% of Schistosoma egg-induced liver granulomas in schistosomiasis hepatic fibrosis (16). M2 macrophages are activated by Th2 cytokines, such as IL-13 and IL-4, which are abundant in chronic infectious. Excessive M2 activation contributes to chronic liver fibrosis by secreting pro-fibrotic cytokines (45, 60). In our cellular model, the M2 macrophage pathway was activated by IL-4/IL-13 cytokines. *In vivo*, a schistosome egg-induced liver fibrosis model was established, and the IL-13/IL-4/Stat6 signalling pathway was stimulated by IL-13, which was released from Schistosoma egg antigens. As shown in Figure 11, IL-4 and IL-13 stimulate macrophage function, leading to the alternative activated M2 phenotype via Stat6 activation. IL-13 and IL-4 activated the IL-4Rα/IL-13Rα1complex, which transduced the signal and activated Stat6, activating transcription of genes typical of M2 polarization, notably Mannose receptor (Mrc1/CD206), PPARγ, Fizz1, and chitinase3-like3 (Chi3l3/Ym1). Meanwhile, Stat6 mediated the alternative activation pathway by up-regulating SOCS1, which suppresses the action of STAT1, a M1 gene. Additionally, the downstream molecules PPARγ and PPARδ regulate a distinct subset of genes that are associated with M2 macrophage activation. Moreover, KLF4 cooperates with Stat6 to promote M2 gene transcription (Mrc1, Fizz1, and PPARγ) and blocks M1 activation by inhibiting the action of nuclear factor kappa-light-chain-enhancer of activated B cell (NF-κB) -p50/p65 (10, 17, 19, 61). Taken together, the above reaction causes an enhanced downstream fibrogenic cytokine response, leading to the generation of hepatic fibrosis.

**Figure 11.**
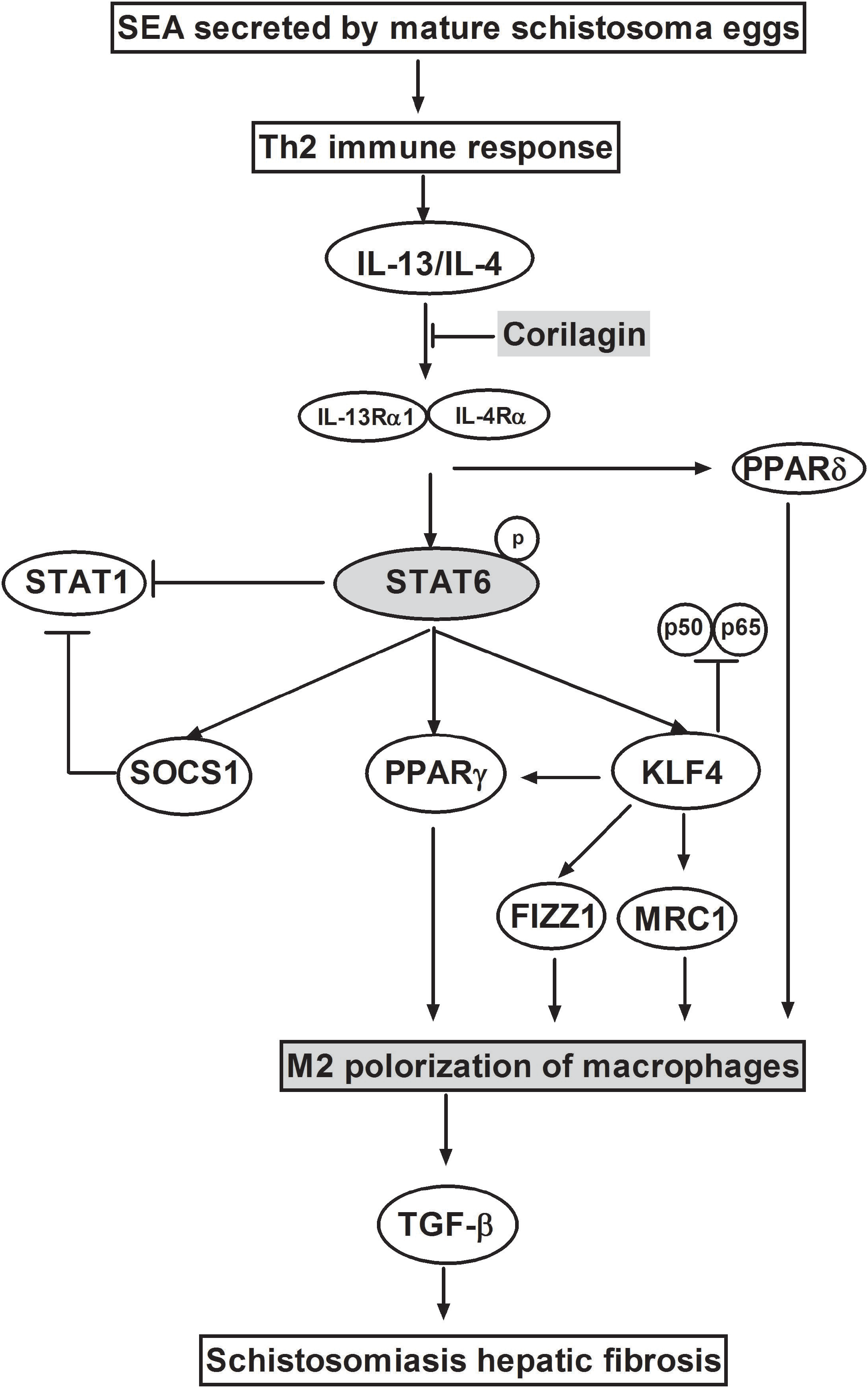
Schematic of the IL-13/IL-4-Stat6 pathway.

Liver fibrosis is a complex chronic disease with multiple causes, such as hepatitis B/C viral infection, steatosis, metabolic disorders, alcohol abuse, Schistosoma infections and autoimmune attacks in liver (62). The causes of hepatic fibrosis are complicated. Developing a comprehensive efficient treatment will not be easy. Meanwhile, side-effects and safety in clinical practice also cannot be overlooked. The good news is that some approaches are currently in preclinical development. For example, siRNA delivery techniques are used as a strategy to target activated myofibroblasts (63). This difficult problem requires more scientific manpower, new technologies and improved quality of research.

In our previous study [31], we focused on exploring whether Corilagin could achieve a regulation on the IL-13-associated pathways in schistosomiasis liver fibrosis. The study was so preliminary that included only mouse model and there was no praziquantel to kill the schistosome. Thereby the effect of Corilagin for schistisomiasis includes inhibition for adult Schistosoma japonicum. The exact effect of Corilagin on egg-induced signaling pathway remained further exploration. Our latest article [64] focuses on membrane protein receptor IL-13α as regulation-up/down target. The main research point for interaction of Corilagin and signaling pathway are also based on IL-13α. In our present study, we chose Stat6 as target for Corilagin interaction. The schistosome were killed before Corilagin treatment and the scope was to investigate the effect on post-parasiticide schistosome egg-induced liver fibrosis via Stat6. The result is another part of effect of Corilagin on schistosomiasis liver fibrosis.

The research investigated the positive inhibitory effect of Corilagin on the progress of schistosome egg-induced liver fibrosis through the IL4/IL13/Stat6 signalling pathways in hepatic alternative activated macrophages *in vitro* and *in vivo*. As a result, the level of Arg1 as well as the mRNA and protein levels of the M2 genes Stat6/P-Stat6, SOCS1, KLF4 and PPAR (γ/δ) were shown to be positively suppressed by Corilagin in IL-13/IL-4-induced cells with or without knockdown/over-expression of Stat6. Meanwhile, in post-parasiticide schistosome egg-induced hepatic fibrosis mice, after treatment with Corilagin, the levels of the M2 markers Fizz1, Ym1, and CD206^+^/CD68^+^ were significantly lowered. There was a similar impact of Corilagin on TGFβ expression, which is one of the strongest factors that leads to liver fibrosis. Corilagin also suppressed the expression of the downstream signalling molecules Stat6/P-Stat6, SOCS1, KLF4 and PPAR (γ/δ) in the IL-4/IL-13/Stat6 pathway in model mice. Corilagin increased mRNA and protein expression of P-P65. Additionally, Corilagin reduced the pathological degree of liver fibrosis.

In conclusion, Corilagin inhibited schistosome egg-induced liver fibrosis via the IL4/IL13/Stat6 signalling pathways in hepatic alternative activated macrophages. These inhibitory effects suggest that Corilagin could be an attractive therapeutic option and supports the need for a more comprehensive study of Corilagin for schistosome egg-induced liver fibrosis treatment. Case in which agents work well in culture or animal models but barely work in patients, or even carry a high risk of side-effects, are not rare. Our follow-up work requires conducting extensive manufacturing validation studies before preclinical development could occur.

## Conflicts of interest

The researchers claim no conflicts of interests.

## Acknowledgement

This study was supported by the National Natural Science Foundation of China (81371840), Hubei Province Health and Family Planning Scientific Research Project (WJ2017Q021), Hubei Provincial Natural Science Foundation of China (2017CFB471), the Fundamental Research Funds for the Central Universities (2017KFYXJJ238), Shandong Provincial Natural Science Foundation of China (2016ZRB14450).

